# Essential role of CFAP53 in sperm flagellum biogenesis

**DOI:** 10.1101/2021.02.04.429673

**Authors:** Bingbing Wu, Xiaochen Yu, Chao Liu, Lina Wang, Tao Huang, Gang Lu, Zi-Jiang Chen, Wei Li, Hongbin Liu

**Author notes:** Corresponding Authors: Dr. Wei Li, Dr. Hongbin Liu. These authors contributed equally to this work (Bingbing Wu, Xiaochen Yu).

## Abstract

The sperm flagellum is essential for male fertility. Despite vigorous research progress towards understanding the pathogenesis of flagellum-related diseases, much remains unknown about the mechanisms underlying the flagellum biogenesis itself. Here, we show that the cilia and flagella associated protein 53 (*Cfap53*) gene is predominantly expressed in testes, and it is essential for sperm flagellum biogenesis. The knockout of this gene resulted in complete infertility in male mice but not in the females. CFAP53 localized to the manchette and sperm tail during spermiogenesis, the knockout of this gene impaired flagellum biogenesis. Furthermore, we identified two manchette and sperm tail-associated proteins that interacted with CFAP53 during spermiogenesis. The disruption of *Cfap53* decreased the expression level of these two proteins and disrupted their localization in spermatids. Together, our results suggest that CFAP53 is an essential protein for sperm flagellum biogenesis, and its mutations might be associated with MMAF.

## Introduction

Infertility is a widespread human health issue, affecting 10%–15% of couples worldwide, and male factors account for around 50% of these cases (Boivin et al., 2007; Tüttelmann et al., 2018). Male infertility is clinically diagnosed as azoospermia, decreased sperm concentration (oligozoospermia), reduced percentage of morphologically normal sperm (teratozoospermia), or lower sperm motility (asthenozoospermia) (Coutton et al., 2015; Ray et al., 2017; Tüttelmann et al., 2018; Zinaman et al., 2000). Spermatozoa are polarized cells composed of two main parts, the head and the flagellum. The flagellum makes up about 90% of the length of the sperm and is essential for sperm motility (Burgess et al., 2003; Mortimer, 2018), and it contains axoneme and peri-axonemal structures, such as the mitochondrial sheath, outer dense fibers, and the fibrous sheath (Mortimer, 2018), and the presence of these structures allows the flagellum to be divided into the connecting piece, midpiece, principal piece, and endpiece (M. S. Lehti & A. Sironen, 2017). Defects in formation of the flagellum disrupt sperm morphology and motility, leading to male infertility (Chemes & Rawe, 2010; Sironen et al., 2020; Turner et al., 2020). Great progress has been made in our understanding of the pathogenesis of flagella-related diseases in recent years, but the pathogenic genes and mechanisms of flagellum biogenesis are far from being fully understood.

The flagellum needs to be integrated with the head in order to function properly during fertilization, and a very complex structure called the sperm head-tail coupling apparatus (HTCA) is necessary for the integration of the sperm head and the flagellum, and defects in this structure result in acephalic spermatozoa syndrome (Wu et al., 2020). Recently, *SUN5, PMFBP1, HOOK1, BRDT, TSGA10*, and *CEP112* have been found to be involved in the assembly of the HTCA, and mutations in these genes are associated with acephalic spermatozoa syndrome (Chen et al., 2018; Li et al., 2017; Sha, Wang, et al., 2020; Sha et al., 2018; Shang et al., 2018; Zhu et al., 2018; Zhu et al., 2016). Abnormalities of the axoneme and accessory structures mainly result in asthenozoospermia, which is associated with morphological flagellar defects such as abnormal tails, irregular mitochondrial sheaths, and irregular residual cytoplasm (Escalier & Touré, 2012; Tu et al., 2020). Previous studies have identified several flagella-associated genes, including *AKAP3, AKAP4, TTC21A, TTC29, FSIP2, DNAH1, DNAH2, DNAH6, DNAH8, DNAH17*, and *DZIP1*, that are involved in sperm flagellum biogenesis (Ben Khelifa et al., 2014; Y. Li et al., 2019; C. Liu et al., 2019; C. Liu et al., 2020; W. Liu et al., 2019; Lv et al., 2020; Martinez et al., 2018; Sha, Wei, et al., 2020; Tu et al., 2019; Turner et al., 2001). Mutations in these genes cause multiple morphological abnormalities of the flagella (MMAF), which is characterized as sperm without flagella or with short, coiled, or otherwise irregular flagella (Ben Khelifa et al., 2014; Touré et al., 2020). There are two evolutionarily conserved bidirectional transport platforms that are involved in sperm flagellum biogenesis, including intramanchette transport (IMT) and intraflagellar transport (IFT) (Kierszenbaum, 2001, 2002; San Agustin et al., 2015). IMT and IFT share similar cytoskeletal components, namely microtubules and F-actin, that provide tracks for the transport of structural proteins to the developing tail (Abraham L. Kierszenbaum et al., 2011), and mutations in *TTC21A, TTC29, SPEF2*, and *CFAP69*, which have been reported to disrupt sperm flagellar protein transport, also lead to MMAF (Dong FN, 2018; C. Liu et al., 2019; Chunyu Liu et al., 2020; W. Liu et al., 2019; Sha et al., 2019).

The cilia and flagella associated protein (CFAP) family, such as *CFAP58, CFAP61, CFAP69, CFAP65, CFAP43, CFAP44, CFAP70*, and *CFAP251*, is associated with flagellum biogenesis and morphogenesis (Beurois et al., 2019; Dong FN, 2018; He et al., 2020; Huang et al., 2020; W. Li et al., 2019; Li et al., 2020; Tang et al., 2017). Previous studies have indicated that the functional role of CFAP53 (also named the coiled-coil domain containing protein CCDC11) is involved in the biogenesis and motility of motile cilia (Narasimhan et al., 2015; Noël et al., 2016; Perles et al., 2012; Silva et al., 2016), and CFAP53 is localized not only to the base of the nodal cilia, but also along the axoneme of the tracheal cilia (Ide et al., 2020). However, the exact localization and function of CFAP53 during spermiogenesis is still poorly understood. In the present study, we used a *Cfap53* knockout mouse model to study the underlying mechanism of CFAP53 in sperm flagellum biogenesis. We demonstrated that CFAP53 is localized to the manchette and the sperm tail of spermatids, and we found that depletion of CFAP53 led to defects in sperm flagellum biogenesis and sperm head shaping. Moreover, we identified two proteins that interacted with CFAP53 during spermiogenesis, namely intraflagellar transport protein 88 (IFT88) and coiled-coil domain containing 42 (CCDC42). *Cfap53* knockout reduced the accumulation of both IFT88 and CCDC42 and disrupted the localization of IFT88 in spermatids. Thus, in addition to uncovering the essential role of CFAP53 in sperm flagellum biogenesis, we also show that CFAP53 might participate in the biogenesis of the sperm flagellum by collaborating with the IMT and IFT pathways.

## Results

### *Cfap53* knockout leads to male infertility

To identify the biological function of CFAP53, we first examined its expression pattern in different tissues and found that it was predominantly expressed in testis (Fig. 1A). Further immunoblotting of mouse testis lysates prepared from different days after birth was carried out. CFAP53 was first detected in testis at postnatal day 7 (P7), and the level increased continuously from postnatal P14 onward, with the highest levels detected in adult testes (Fig. 1B). This time course corresponded with the onset of meiosis, suggesting that CFAP53 might have an essential role in spermatogenesis.

**Fig. 1.**
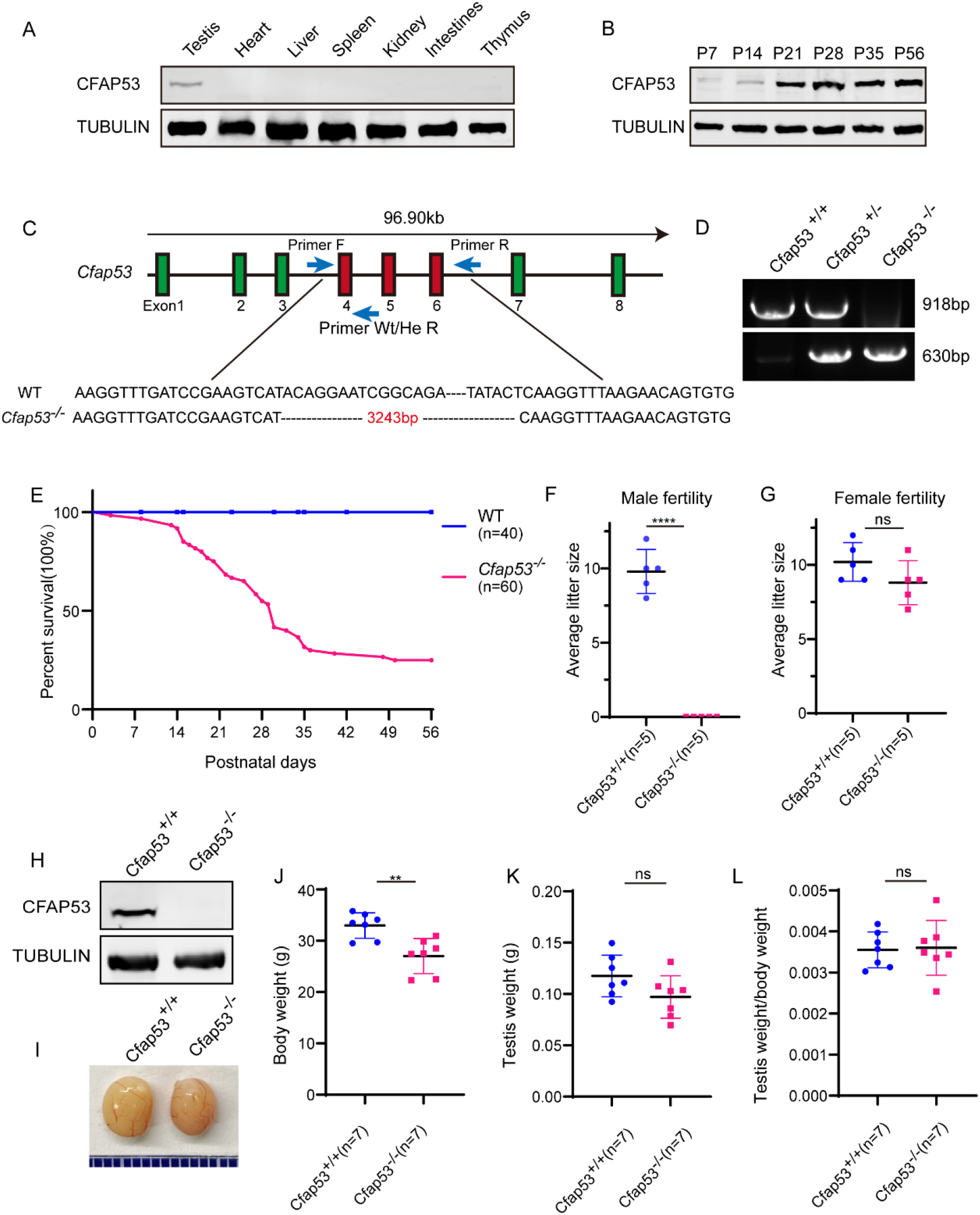
The generation of *Cfap53* knockout mice. (A) CFAP53 was predominately expressed in testis. Immunoblotting of CFAP53 was performed in testis, heart, liver, spleen, kidney, intestines, and thymus with Tubulin serving as the control. (B) CFAP53 was expressed starting in P7 testes. Tubulin served as the control. (C) The generation of *Cfap53*^−*/*–^ mice lacking exons 4 to 6. (D) Genotyping of *Cfap53 Cfap53*^−*/*–^ mice. (E) Survival rate of postnatal *Cfap53*^−*/*–^ mice (n = 60). (F) The average litter size of *Cfap53*^*+/+*^ and *Cfap53*^−*/*–^ male mice at 3 months (n = 5 independent experiments). *Cfap53*^−*/*–^ male mice were completely sterile. Data are presented as the mean ± SD. \*\*\*\**P* < 0.0001. (G) The average litter size of *Cfap53*^*+/+*^ and *Cfap53*^−*/*–^ female mice at 3 months (n = 5 independent experiments). *Cfap53*^−*/*–^ female mice were fertile. Data are presented as the mean ± SD. (H) Immunoblotting of CFAP53 in *Cfap53*^*+/+*^ and *Cfap53*^−*/*–^ testes. Tubulin served as the control. (I) The testis sizes of *Cfap53*^*+/+*^ and *Cfap53*^−*/*–^ mice were similar to each other. Data are presented as the mean ± SD. (J) The body weights of *Cfap53*^−*/*–^ male mice were lower compared to *Cfap53*^*+/+*^ male mice (n = 7 independent experiments). Data are presented as the mean ± SD. \*\**P* < 0.01. (K) The testis weights of *Cfap53*^*+/+*^ and *Cfap53*^−*/*–^ male mice (n = 7 independent experiments). Data are presented as the mean ± SD. (L) The ratio of testis weight/body weight in *Cfap53*^*+/+*^ and *Cfap53*^−*/*–^ male mice (n = 7 independent experiments). Data are presented as the mean ± SD.

To characterize the potential functions of CFAP53 during spermatogenesis, *Cfap53*^−*/*–^ mice were created using the CRISPR-Cas9 system from Cyagen Biosciences. Exon 4 to exon 6 of the *Cfap53* gene was selected as the target site (Fig. 1C). The founder animals were genotyped by genomic DNA sequencing and further confirmed by polymerase chain reaction – the total size of the *Cfap53* locus in *Cfap53*^*+/+*^ mice was 918 bp, while the size of the locus in *Cfap53*^−*/*–^ mice was 630 bp (Fig. 1D). Immunoblotting of testis indicated that the CFAP53 protein was successfully eliminated in *Cfap53*^−*/*–^ mice (Fig. 1H). Because we cannot obtain adult homozygous *Cfap53*^−*/*–^ mice in the C57BL/6J (C57) background, the heterozygous *Cfap53* mutated mice in the C57 background were further crossed with wild-type (WT) ICR mice, and the resulting heterozygotes were interbred to obtain homozygous *Cfap53*^−*/*–^ mice in the C57/ICR background. The offspring genotypes deviated from Mendelian ratios (115:237:60 for *Cfap53*^*+/+*^: *Cfap53*^*+/-*^: *Cfap53*^−*/*–^), suggesting an increased prenatal lethal rate in *Cfap53*^−*/*–^ mice. We analyzed 25 *Cfap53*^−*/*–^ mice, and 48% (12/25) of them presented with situs inversus totalis (SIT) and nearly 8% (2/25) had situs inversus abdominalis (SIA). In addition, 32% (8/25) of the *Cfap53*^−*/*–^ mice developed hydrocephalus (Supplementary Fig. 1).

A total of 72% (43/60) of the *Cfap53*^−*/*–^ mice that survived after birth died within 6 weeks, while 25% (15/60) of the *Cfap53*^−*/*–^ mice lived longer than 8 weeks of age (Fig. 1E). We further examined the fertility of *Cfap53* male and female knockout mice. *Cfap53* male knockout mice exhibited normal mounting behaviors and produced coital plugs, but *Cfap53*^−*/*–^ male mice failed to produce any offspring after mating with WT adult female mice. In contrast, *Cfap53*^−*/*–^ female mice generated offspring after mating with WT males (Fig. 1F–G). Thus, the disruption in *Cfap53* resulted in male infertility but did not affect the fertility of *Cfap53*^−*/*–^ female mice.

### The knockout of *Cfap53* results in MMAF

To further investigate the cause of male infertility, we first observed the adult *Cfap53*^−*/*–^ testis structure at both the gross and histological levels. The body weight of *Cfap53*^−*/*–^ male mice was reduced compared to *Cfap53*^*+/+*^ male mice (Fig. 1J), while there were no significant differences in the testis size, testis weight, or testis/body weight ratio between *Cfap53*^−*/*–^ and *Cfap53*^*+/+*^ male mice (Fig. 1I, K, L). We then observed the transverse sections of the *Cfap53*^−*/*–^ cauda epididymis by hematoxylin and eosin (H&E) staining and found that there was a complete lack of spermatozoa or only a few spermatozoa in the epididymal lumen of *Cfap53*^−*/*–^ mice (Fig. 2A, red arrowhead). We examined the spermatozoa released from the caudal epididymis and found the sperm count in the *Cfap53*^−*/*–^ mice to be significantly decreased compared with WT mice (Fig. 2B). To determine the morphological characteristics of the spermatozoa, we performed single-sperm immunofluorescence of lectin peanut agglutinin (PNA), which is used to visualize the acrosomes of spermatozoa. The *Cfap53*^−*/*–^ caudal epididymis only contained malformed spermatozoa exhibiting the prominent MMAF phenotype of short, coiled, or absent flagella compared with *Cfap53*^*+/+*^ mice. In addition to the flagella abnormality, *Cfap53*^−*/*–^ mice had abnormal sperm heads (Fig. 2C). The ratio of spermatozoa with abnormal heads and flagella is shown in Fig. 2D. Abnormal sperm head with short tail and normal sperm head with curly tail were the major defect categories. Immunofluorescence analysis with MitoTracker, which is used to visualize mitochondria, showed that the mitochondrial sheath was malformed in the *Cfap53*^−*/*–^ spermatozoa (Supplementary Fig. 2).

**Fig. 2.**
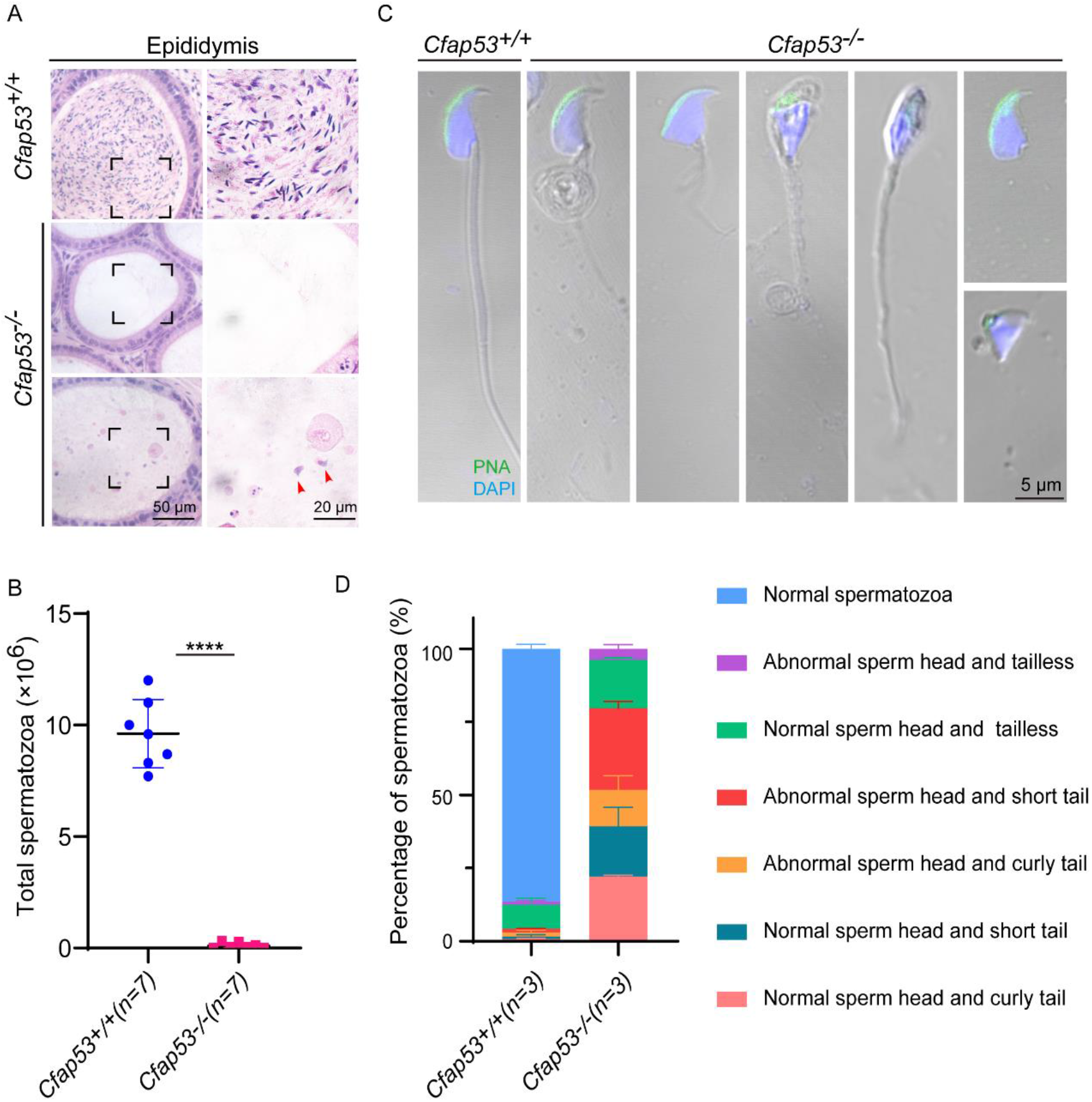
CFAP53 knockout results in MMAF. (A) H&E staining of the caudal epididymis from *Cfap53*^*+/+*^ and *Cfap53*^−*/*–^ male mice. (B) The sperm counts in the caudal epididymis were significantly decreased in the *Cfap53*^−*/*–^ male mice (n = 7 independent experiments). Data are presented as the mean ± SD. \*\*\*\**P* < 0.0001. (C) Immunofluorescence staining of PNA in Cfa*p53*^*+/+*^ and *Cfap53*^−*/*–^ spermatozoa, indicating abnormal spermatozoa such as abnormal head and coiled, short, or absent flagella. (D) Quantification of different categories of abnormal spermatozoa (n = 3 independent experiments). Data are presented as the mean ± SD. The statistical significance of the differences between the mean values for the different genotypes was measured by Student’s t-test with a paired 2-tailed distribution.

### CFAP53 is required for spermatogenesis

To address the question of why *Cfap53* knockout results in MMAF, we conducted Periodic acid–Schiff (PAS) staining to determine the stages of spermatogenesis in *Cfap53*^−*/*–^ and WT testes. The most prominent defects were observed in the spermatids at the stages of spermatogenesis, where abnormally elongated and constricted sperm head shapes were identified (Fig. 3A, asterisks). In addition, some dead cells could be detected in the *Cfap53*^−*/*–^ seminiferous tubules (Fig. 3A). To clarify the detailed morphological effects of the *Cfap53* mutation on the structure of sperm heads, we analyzed the process of sperm head shaping between *Cfap53*^−*/*–^ and *Cfap53*^*+/+*^ mice. Notably, from step 1 to step 8 the acrosome and nucleus morphology in *Cfap53*^−*/*–^ spermatids was normal compared with *Cfap53*^*+/+*^ spermatids. Head shaping started at step 9 to step 10, and the morphology of the elongated *Cfap53*^−*/*–^ spermatid heads was normal compared with that of *Cfap53*^*+/+*^ mice, whereas abnormal club-shaped heads (Fig. 3B) were seen in step 11 spermatids in *Cfap53*^−*/*–^ mice. This phenomenon became more apparent between step 11 and step 16 (Fig. 3B). Taken together, these results indicate that CFAP53 is required for normal spermatogenesis.

**Fig. 3.**
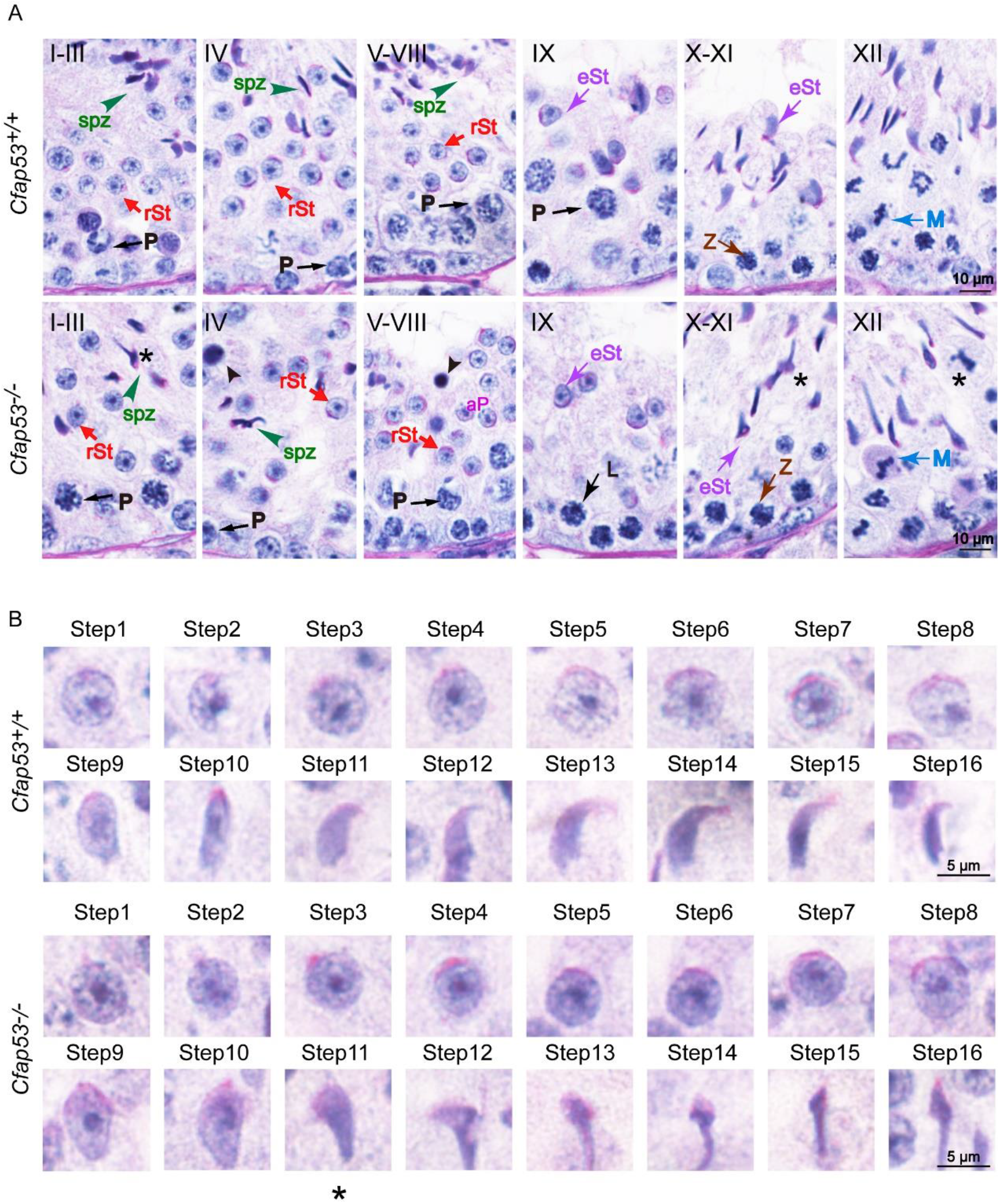
Spermatogenesis defects of *Cfap53*^−*/*–^ mice. (A) PAS staining of testes sections from *Cfap53*^*+/+*^ and *Cfap53*^−*/*–^ mice. Defects in the nuclear shape of several elongating spermatids were clearly evident in the *Cfap53*^−*/*–^ seminiferous tubule (asterisks). Apoptotic bodies were detected in *Cfap53*^−*/*–^ testes sections (black arrowheads). P: pachytene spermatocyte, L: leptotene spermatocyte, Z: zygotene spermatocyte, M: meiotic spermatocyte, rST: round spermatid, eST: elongating spermatid, spz: spermatozoa. (B) The PAS staining of spermatids at different steps from *Cfap53*^*+/+*^ and *Cfap53*^−*/*–^ mice. From step 1 to step 10 spermatids, the head morphology was roughly normal in *Cfap53*^−*/*–^ mice. Abnormal, club-shaped heads (asterisk) were first seen in step 11 spermatids in *Cfap53*^−*/*–^ mice.

H&E staining was used to further observe the morphological changes of the seminiferous tubules. The seminiferous tubules of *Cfap53*^*+/+*^ mice had a tubular lumen with flagella appearing from the developing spermatids. In contrast, the flagella were absent in the seminiferous tubules of *Cfap53*^−*/*–^ mice (Fig. 4A). Immunofluorescence staining for α/β-tubulin, the specific flagellum marker, further confirmed the defects in flagellum biogenesis resulting from the knockout of *Cfap53* (Fig. 4B). These observations clearly suggest that CFAP53 plays an important role in flagellum biogenesis.

**Fig. 4.**
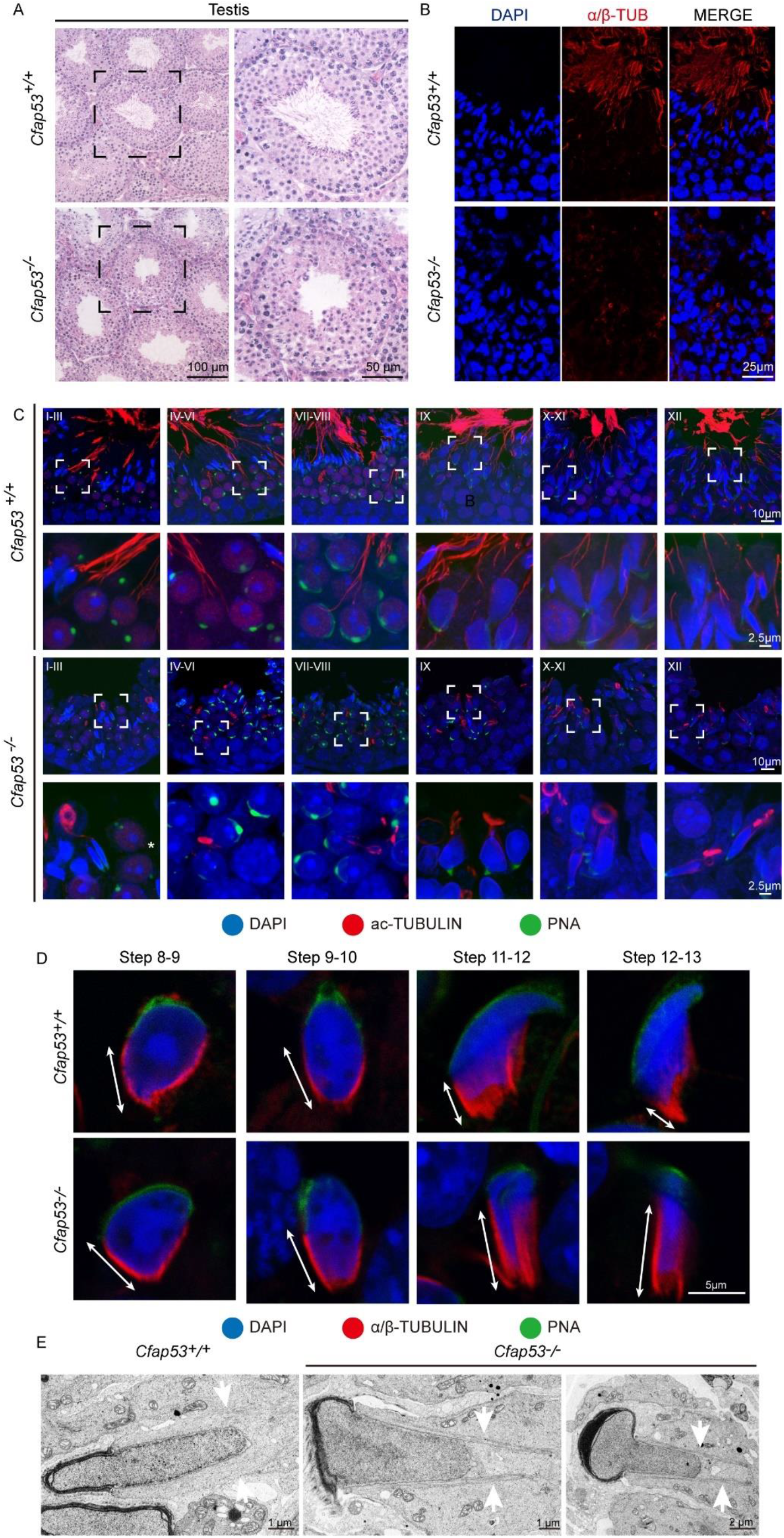
Sperm flagellum biogenesis defects and abnormal manchettes in *Cfap53*^−*/*–^ mice. (A) H&E staining of testes sections from *Cfap53*^*+/+*^ and *Cfap53*^−*/*–^ male mice. (B) Immunofluorescence of anti-α/β-tubulin (red) antibodies in testes sections from *Cfap53*^−*/*–^ male mice show flagellar defects. (C) Comparison of flagellum biogenesis in testes sections from *Cfap53*^*+/+*^ and *Cfap53*^−*/*–^ mice at different stages. Sperm flagella were stained with acetylated Tubulin (red), the acrosome was stained with PNA lectin histochemistry (green), and the nucleus was stained with DAPI (blue). Flagellum formation was first observed at stages I-III of the seminiferous epithelial cycle in *Cfap53*^*+/+*^ mice, while sperm tails were not detected (asterisks) at stages I-III in testes sections in *Cfap53*^−*/*–^ mice. From stages IV–VI, sperm flagellum biogenesis defects were clearly seen in *Cfap53*^−*/*–^ testis sections. (D) Comparison of manchette formation between *Cfap53*^*+/+*^ and *Cfap53*^−*/*–^ spermatids at different steps. The manchette was stained with α/β-tubulin (red), the acrosome was stained with PNA lectin histochemistry (green), and the nucleus was stained with DAPI (blue). The distance from the perinuclear ring to the caudal side of the nucleus is indicated by white arrows. During steps 12 and 13, the distance was reduced in *Cfap53*^*+/+*^ spermatids, while the manchette of *Cfap53*^−*/*–^ spermatids displayed abnormal elongation. (E) Transmission electron microscope images of *Cfap53*^−*/*–^ step 11–13 spermatids showing the perinuclear ring constricting the sperm nucleus and causing abnormal sperm head formation. White arrows indicate the manchette microtubules.

### CFAP53 is required for sperm flagellum biogenesis and correct manchette function

In order to determine the causes of the abnormal sperm morphology in *Cfap53*^−*/*–^ mice, we investigated the effect of *Cfap53* knockout on flagellum biogenesis using the antibody against acetylated tubulin, a flagellum-specific marker. Unlike the well-defined flagellum of the control group, the axoneme was absent in step 2–3 spermatids in *Cfap53*^−*/*–^ mice (Fig. 4C, asterisks). In steps 4–6, abnormally formed flagella were seen in *Cfap53*^−*/*–^ testis sections (Fig. 4C). The presence of long and abnormal spermatid heads suggested defects in the function of the manchette, which is involved in sperm head shaping. Immunofluorescence staining for α/β-tubulin antibody showed that manchette formation was normal in step 8 to step 10 spermatids in *Cfap53*^−*/*–^ mice, while step 11 to step 13 spermatids of *Cfap53*^−*/*–^ mice had abnormally long manchettes compared with WT controls. (Fig. 4D). We performed transmission electron microscopy to study the organization of the sperm manchette in detail in *Cfap53*^−*/*–^ mice. During the chromatin condensation period starting from step 11 spermatids, the manchette of *Cfap53*^− */*–^ mice appeared abnormally long and the perinuclear ring constricted the sperm nucleus, causing severe defects in sperm head formation (Fig. 4E). Thus, deletion of *Cfap53* causes severe defects in sperm flagellum biogenesis and manchette function.

### CFAP53 localizes to the manchette and the sperm tail

In order to determine the functional role of CFAP53 during spermiogenesis, we investigated the subcellular localization of CFAP53 during spermatogenesis in mice using an anti-CFAP53 antibody. The CFAP53 signal was first observed as two adjacent dots nearby the nucleus in spermatocytes and early round spermatids (Fig. 5A), and these results were consistent with the protein expression patterns (Fig. 1B). During the elongation of the spermatids (step 9 to step 14), CFAP53 could be detected as a skirt-like structure that encircled the elongating spermatid head, and the protein was subsequently located to the sperm tail around step 14 to step 15. Compared to *Cfap53*^*+/+*^ mice, there was no CFAP53 staining detected in the germ cells of *Cfap53*^−*/*–^ male mice (Fig. 5A). To determine whether CFAP53 associates with microtubular structures, the localization of CFAP53 in the elongating and elongated spermatid was subsequently co-stained with antibodies against α-tubulin (a manchette marker) and against CFAP53. In the elongating spermatid CFAP53 colocalized with the manchette microtubules. CFAP53 was further identified at the sperm tail, whereas α-tubulin marked the whole tail in the elongated spermatids (Fig. 5B). Taken together, these results indicate that CFAP53 might participate in manchette formation and flagellum biogenesis.

**Fig. 5.**
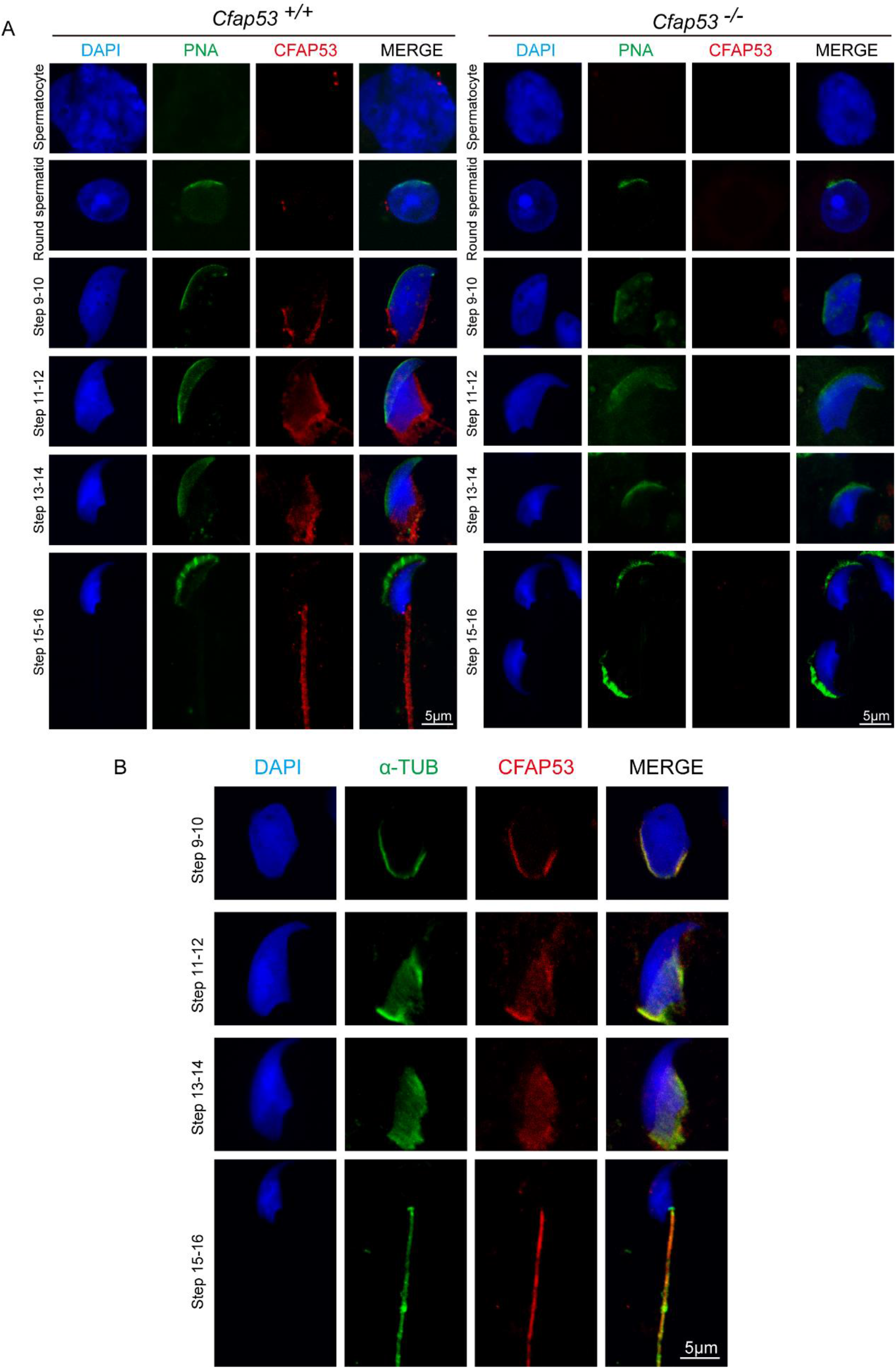
Localization of CFAP53 in developing germ cells. (A) Testicular germ cells were prepared from *Cfap53*^*+/+*^ and *Cfap53*^−*/*–^ adult mouse testis, and immunofluorescence staining was performed with antibodies to CFAP53 (red). The acrosome was stained with PNA lectin histochemistry (green), and the nucleus was stained with DAPI (blue). (B) Testicular spermatids of WT adult mouse testes were stained with antibodies against α-tubulin (green) and CFAP53 (red). The nucleus was stained with DAPI (blue). In step 9–14 spermatids, CFAP53 was detected at the manchette. In 15–16 spermatids, CFAP53 was located at the sperm tail.

### CFAP53 interacts with IFT88 and CCDC42

Sperm flagellum biogenesis requires protein delivery to the assembly sites via IMT and IFT (Lehti & Sironen, 2016). According to the localization and functional role of CFAP53 during spermiogenesis, some IFT/IMT-related genes were chosen to determine their interactions with CFAP53 by coimmunoprecipitation (co-IP) assays. IFT88 and IFT20, which belong to the IFT family, are involved in protein transport, and their depletion affects sperm flagellum biogenesis (A. L. Kierszenbaum et al., 2011; Zhang et al., 2016). CCDC42 is involved in IMT and is essential for sperm flagellum biogenesis (Pasek et al., 2016; Tapia Contreras & Hoyer-Fender, 2019). To determine their relationship with CFAP53, HEK293T cells were co-transfected with MYC-tagged CFAP53 and IFT88-GFP, CCDC42-GFP, or GFP-tagged empty vector plasmid as a control, and we found that both IFT88 and CCDC42 were coimmunoprecipitated with CFAP53-MYC (Fig. 6A-C). We did not detect any interaction between IFT20-FLAG and CFAP53-MYC using the same strategy (Fig. 6D). To further determine their relationship, we performed immunoblotting of IFT88 and CCDC42 in the testes of *Cfap53*^*+/+*^ and *Cfap53*^−*/*–^ mice, and we found that the expression levels of IFT88 and CCDC42 were significantly reduced in the testes of *Cfap53*^−*/*–^ mice (Fig. 6E, F). Taken together, we show that CFAP53 interacts with IFT88 and CCDC42 and that the knockout of *Cfap53* decreases the expression of CCDC42 and IFT88.

**Fig. 6.**
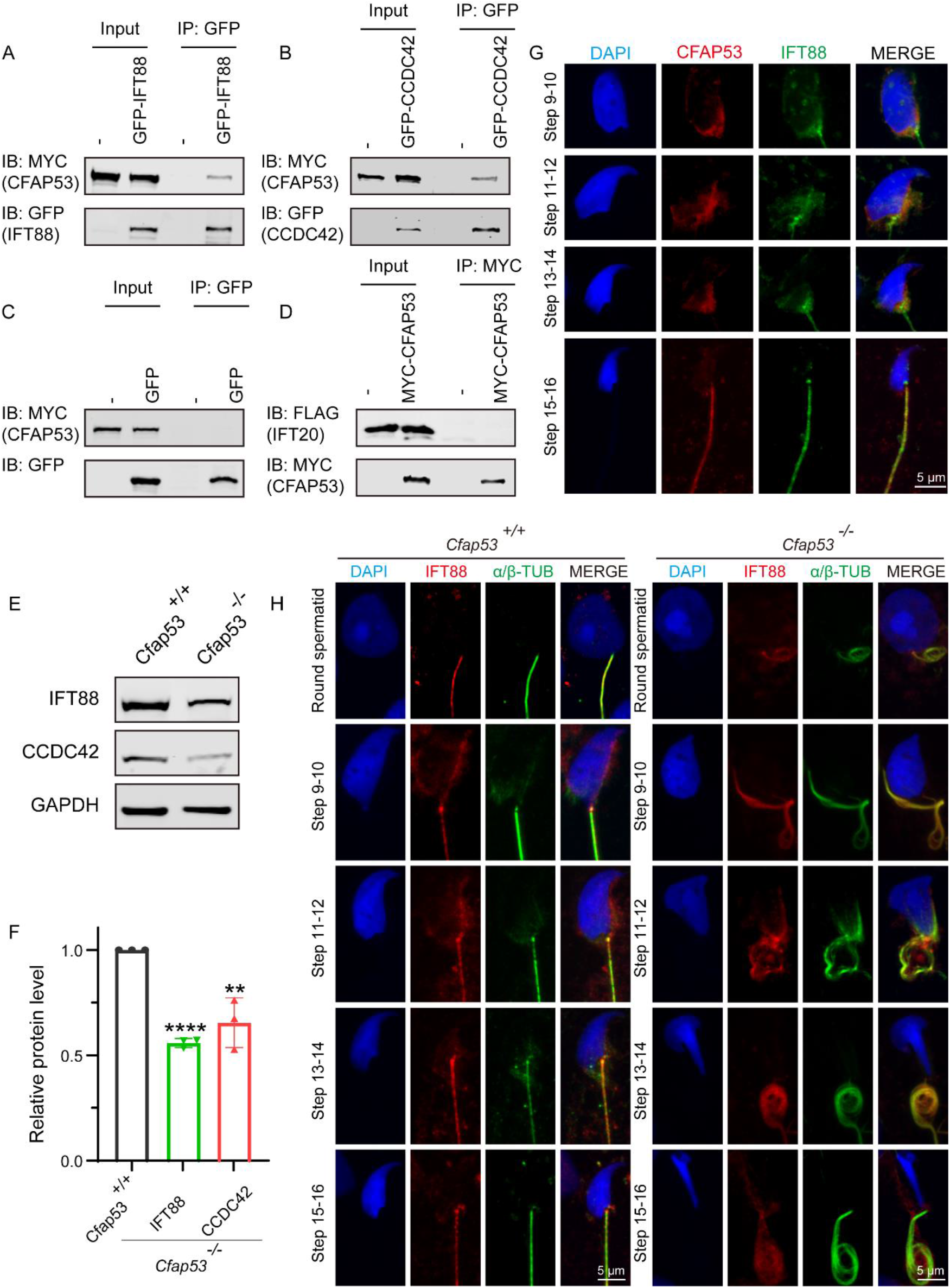
CFAP53 interacts with IFT88 and CCDC42. (A) and (B) CFAP53 interacts with IFT88 and CCDC42. *pCS2-Myc-Cfap53* and *pEGFP-GFP-Ift88* or *pEGFP-GFP-Ccdc42* were co-transfected into HEK293T cells. At 48 h after transfection, the cells were collected for immunoprecipitation (IP) with anti-GFP antibody and analyzed with anti-MYC and anti-GFP antibodies. (C) *pCS2-Myc-Cfap53* and Empty vector were co-transfected into HEK293T cells. At 48 h after transfection, the cells were collected for IP with anti-GFP antibody and analyzed with anti-MYC and anti-GFP antibodies. (D) *pCS2-Myc-Cfap53* and *pRK-Flag-Ift20* were co-transfected into HEK293T cells. At 48 h after transfection, the cells were collected for IP with anti-MYC antibody and analyzed with anti-MYC and anti-FLAG antibodies. (E) Western blot analysis showing IFT88 and CCDC42 protein levels in *Cfap53*^*+/+*^ and *Cfap53*^−*/*–^ mouse testis lysates. GAPDH served as the loading control. (F) Quantification of the relative protein levels of IFT88 and CCDC42 using the Odyssey software and compared with the control group (n = 3 independent experiments). Data are presented as the mean ± SD. The statistical significance of the differences between the mean values for the different genotypes was measured by Student’s t-test with a paired, 2-tailed distribution. \*\**P* < 0.01 and \*\*\*\**P* < 0.0001. (G) The immunofluorescence analysis of IFT88 (green) and CFAP53 (red) was performed in testicular germ cells. The nucleus was stained with DAPI (blue). (H) Immunofluorescence staining with antibodies against IFT88 (red) and α/β-tubulin (green) in spermatids at different developmental stages from *Cfap53*^*+/+*^ and *Cfap53*^−*/*–^ adult mice. The nucleus was stained with DAPI (blue).

Both IFT88 and CCDC42 localize to the manchette, the sperm connecting piece, and the sperm tail during spermatogenesis (A. L. Kierszenbaum et al., 2011; Pasek et al., 2016; Tapia Contreras & Hoyer-Fender, 2019). Because the antibody against CCDC42 does not function for immunofluorescence, we focused on IFT88. Similar to the knockout of *Cfap53*, spermatozoa in the *Ift88* mutant mouse had absent, short, or irregular tails with malformed sperm heads (A. L. Kierszenbaum et al., 2011; San Agustin et al., 2015). In order to further investigate the effect of *Cfap53* knockout on IFT88 localization and its potential interaction with CFAP53 during sperm development, we co-stained the differentiating spermatids with antibodies against IFT88 and CFAP53. We found that IFT88 located to the manchette, the HTCA, and the sperm tail as previously reported (A. L. Kierszenbaum et al., 2011; San Agustin et al., 2015). CFAP53 co-localized with IFT88 in the manchette of the elongating spermatid and in the sperm tail of the elongated spermatid (Fig. 6G). We next detected IFT88 localization in the different steps of spermatid development in *Cfap53*^*+/+*^ and *Cfap53*^−*/*–^ mice. The IFT88 signal was first observed in the tail of round spermatids and continued to be detected in the elongated spermatid. Unlike the well-defined flagellum signal of the control group, the detectable IFT88 signal was abnormal in the spermatids of *Cfap53*^−*/*–^ mice (Fig. 6H). This result suggests that both CFAP53 and IFT88 might cooperatively participate in flagellum biogenesis during spermiogenesis.

## Discussion

In this study, we have identified the essential role of CFAP53 in spermatogenesis and male fertility by generating *Cfap53*^−*/*–^ mice with the deletion of exons 4–6. Sperm flagellum biogenesis begins in early round spermatids, where the axoneme extends from the distal centriole (Mari S Lehti & Anu Sironen, 2017). It has been reported that the expression of many genes that are necessary for sperm flagellum biogenesis is significantly increased at approximately 12 days after birth (Horowitz et al., 2005), and we found that CFAP53 expression was upregulated in the testes between 7 and 14 days after birth (Fig. 1B), which was consistent with the timing of axoneme formation. Previous studies have shown that CFAP53 is located at the basal body and on centriolar satellites in retinal pigment epithelial cells and on the ciliary axonemes in zebrafish kidneys and human respiratory cells (Narasimhan et al., 2015; Silva et al., 2016). In mice, CFAP53 is located at the base of the nodal cilia and the tracheal cilia, as well as along the axonemes of the tracheal cilia (Ide et al., 2020). Depletion of CFAP53 disrupts the subcellular organization of satellite proteins and lead to primary cilium assembly abnormalities (Silva et al., 2016), and knockout of CFAP53 disrupts ciliogenesis in human tracheal epithelial multiciliated cells, in *Xenopus* epidermal multiciliated cells, and in zebrafish Kupffer’s vesicle and pronephros (Narasimhan et al., 2015; Noël et al., 2016; Silva et al., 2016). The mammalian sperm flagellum contains an axoneme composed of a 9+2 microtubule arrangement, which is similar to that of motile cilia. Our study further showed that axoneme formation was impaired in early round spermatids in *Cfap53*^−*/*–^ mice (Fig. 4C), thus demonstrating that CFAP53 is essential for sperm flagellum biogenesis.

We found that CFAP53 localized on the manchette and tail during spermiogenesis, and it could interact with IFT88 and CCDC42, both of which colocalized to the same positions during spermiogenesis (Fig. 6A, B, G). In addition, it has been reported that CFAP53 also interacts with KIF3A in adult testis (Lehti et al., 2013). All of these partner proteins are related to manchette and flagellum biogenesis, with the manchette being one of the transient skirt-like microtubular structures that are required for the formation of sperm flagella and the shaping of the head during spermatid elongation (Kierszenbaum & Tres, 2004; Lehti & Sironen, 2016). It has been proposed that flagellar structure proteins and motor proteins are transported through the manchette via IMT to the base of the sperm flagellum and via IFT to the developing sperm flagellum (Abraham L. Kierszenbaum et al., 2011; Lehti & Sironen, 2016). Both IMT and IFT provide the bidirectional movement of multicomponent transport systems powered by molecular motors along the microtubules, and both are essential for axoneme assembly (Mari S Lehti & Anu Sironen, 2017). Molecular motors (kinesin-2 and dynein 2) move cargo proteins associated with protein rafts consisting of IFT proteins (Chien et al., 2017; Kierszenbaum, 2002; Zhu et al., 2020), and KIF3A, the motor subunit of kinesin-2, works as an anterograde motor for transporting IFT complex B during the development of the sperm tail (Lehti et al., 2013; Marszalek JR, 1999). KIF3A localizes to the manchette, the basal body, and the axoneme of spermatids, and disruption of KIF3A affects the formation of the manchette and further disrupts the delivery of proteins to the sperm tail (Lehti et al., 2013). IFT88 is an IFT complex B protein that is regarded as a member of the IMT machinery. Notably, the reproductive phenotype of *Ift88* knockout male mice was similar to what we observed in *Cfap53* knockout mice (San Agustin et al., 2015). Thus, CFAP53 might function in collaboration with IFT88 and KIF3A in flagellum biogenesis via IMT and IFT (Fig. 7).

**Fig. 7.**
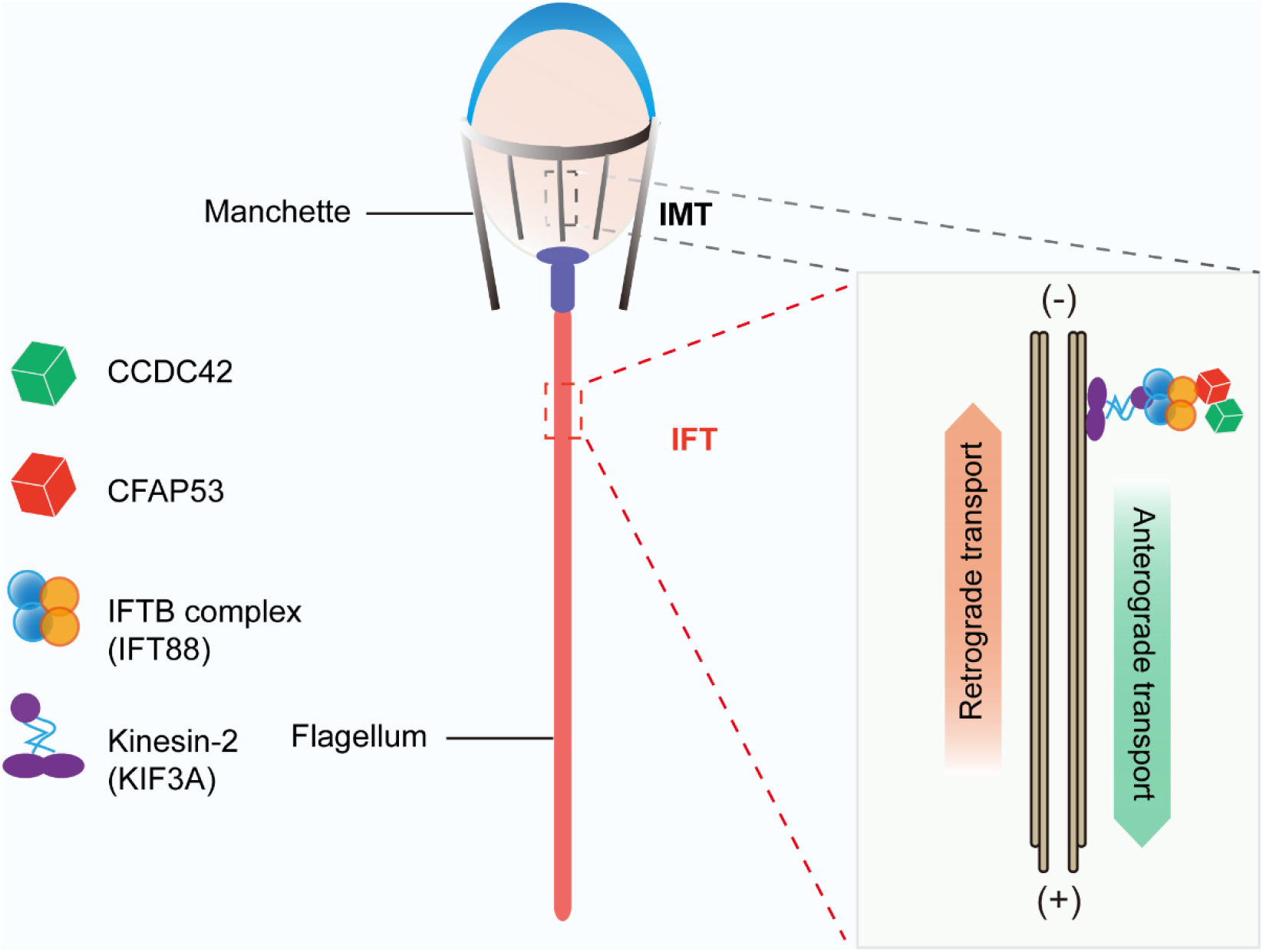
Proposed model for the functional role of CFAP53 during flagellum biogenesis. CFAP53 works as a cargo protein, and it is involved in the stabilization of other cargo proteins, such as CCDC42, that need to be transported to the developing sperm tail. During transport, CFAP53 interacts with IFT complex B member IFT88 and kinesin-2 motor subunit KIF3A for targeting to its destination.

Previous studies have shown that *Ccdc42* knockout male mice have abnormal head shapes and axoneme assembly defects, and it was speculated that CCDC42 might be a passenger protein transported via the manchette towards the developing tail (Pasek et al., 2016; Tapia Contreras & Hoyer-Fender, 2019). As a partner protein of CCDC42 and IFT88, once CFAP53 was depleted the expression levels of these two partner proteins were also decreased significantly (Fig 6E, F). These results raise the question of whether CFAP53 works as a cargo protein or as a component of the IFT and IMT machineries. Given that CFAP53 also localized on sperm flagella, we are drawn to the conclusion that CFPA53 works as a cargo protein and that and it might be involved in the stabilization of other cargo proteins, such as CCDC42, that are transported by IFT and IMT (Fig 7). As for the decreasing of IFT88, this is likely caused by the decreasing demand for cargo protein transportation via IMT and IFT to the developing sperm tail. Accordingly, our current conclusions can be further expanded to other flagellum-associated proteins, and the depletion of some cargo proteins that need to be transported by either IMT or IFT should have similar phenotypes due to flagellum biogenesis defects, and at least some of the MMAF might be caused by this mechanism.

## Methods

### Animals

The mouse *Cfap53* gene (Transcript: ENSMUSG00000035394) is 96.90 kb and contains 8 exons and is located on chromosome 18. Exon 4 to exon 6 was chosen as the target site, and *Cfap53*^−*/*–^ mice were generated using the CRISPR-Cas9 system from Cyagen Biosciences. The gRNA and Cas9 mRNA were co-injected into fertilized eggs of C57BL/6 mice to generate a targeted line with a 3243 bp base deletion, AAG GTT TGA TCC GAA GTC AT – 3243 bp – CAA GGT TTA AGA ACA GTG TG. The founder animals were genotyped by genomic DNA sequencing. For *Cfap53*^−*/*–^ mice, the specific primers were Forward: 5’-GAG GGA ATA GGT TTC TGG GTA GGT G-3’ and Reverse: 5’-ACC CTT CTG GTC CCT CAG TCA TCT-3’, yielding a 630 bp fragment. For *Cfap53* wild-type mice, the specific primers were Forward: 5’-GAG GGA ATA GGT TTC TGG GTA GG TG-3’ and Reverse: 5’-AGC AGC AGT GAA ACT TCA AAC ATG G-3’, yielding a 918 bp fragment. All of the animal experiments were performed according to approved institutional animal care and use committee (IACUC) protocols (#08-133) of the Institute of Zoology, Chinese Academy of Sciences.

### Plasmids

Mouse *Cfap53* was obtained from mouse testis cDNA and cloned into the pCMV-Myc vector using the Clon Express Ultra One Step Cloning Kit (C115, Vazyme). Mouse *Ccdc42* and *Ift88* were obtained from mouse testis cDNA and cloned into the pEGFP-C1 vector using the Clon Express Ultra One Step Cloning Kit (C115, Vazyme). Mouse *Ift20* was obtained from mouse testis cDNA and cloned into the pRK vector using the Clon Express Ultra One Step Cloning Kit (C115, Vazyme).

### Antibodies

Mouse anti-CFAP53 polyclonal antibody (aa 216–358) was generated by Quan Biotech (Wuhan, China) and was used at a 1:10 dilution for immunofluorescence and 1:200 dilution for western blotting. Mouse anti-GFP antibody (1:1000 dilution, M20004L, Abmart, Shanghai, China), rabbit anti-MYC antibody (1:1000 dilution, BE2011, EASYBIO, Beijing, China), mouse anti-FLAG antibody (1:1000 dilution, F1804, Sigma, Shanghai, China), and rabbit anti-CCDC42 antibody (1:500 dilution, abin2785068, antibodies-online, Beijing, China) were used for western blotting. Rabbit anti-α-tubulin antibody (AC007, ABclonal, Wuhan, China) was used at a 1:100 dilution for immunofluorescence and at a 1:10000 dilution for western blotting. IFT88 polyclonal antibody (13967-1-AP, Proteintech, Wuhan, China) was used at a 1:50 dilution for immunofluorescence and at a 1:1000 dilution for western blotting. Mouse anti-α/β-tubulin antibody (1:100, ab44928, Abcam, Shanghai, China) and mouse anti-acetylated-a-tubulin antibody (T7451, 1:1000 dilution, Sigma) were used for immunofluorescence. The secondary antibodies were goat anti-rabbit FITC (1:200, ZF-0311, Zhong Shan Jin Qiao, Beijing, China), goat anti-rabbit TRITC (1:200 dilution, ZF-0316, Zhong Shan Jin Qiao), goat anti-mouse FITC (1:200 dilution, ZF-0312, Zhong Shan Jin Qiao), and goat anti-mouse TRITC (1:200 dilution, ZF-0313, Zhong Shan Jin Qiao). The Alexa Fluor 488 conjugate of lectin PNA (1:400 dilution, L21409, Thermo Fisher Scientific, Shanghai, China) and MitoTracker Deep Red 633 (1:1500 dilution, M22426, Thermo Fisher Scientific) were used for immunofluorescence.

### Immunoprecipitation

Transfected HEK293T cells were lysed in ELB buffer (50 mM HEPES, 250 mM NaCl, 0.1% NP-40, 1 mM PMSF, and complete EDTA-free protease inhibitor cocktail (Roche)) for 30 min on ice and centrifuged at 12000 × *g* for 15 min. For immunoprecipitation, cell lysates were incubated with anti-GFP antibody overnight at 4°C and then incubated with protein A-Sepharose (GE, 17-1279-03) for 3 hours at 4°C Thereafter, the precipitants were washed four times with ELB buffer, and the immune complexes were eluted with sample buffer containing 1% SDS for 10 min at 95°C and analyzed by immunoblotting.

### Immunoblotting

Proteins obtained from lysates or immunoprecipitates were separated by SDS-PAGE and electrotransferred onto a nitrocellulose membrane. The membrane was blocked in 5% skim milk (BD, 232100) and then incubated with corresponding primary antibodies and detected by Alexa Fluor 680 or 800-conjugated goat anti-mouse or Alexa Fluor 680 or 800-conjugated goat anti-rabbit secondary antibodies. Finally, they were scanned using the ODYSSEY Sa Infrared Imaging System (LI-COR Biosciences, Lincoln, NE, RRID:SCR_014579).

### Mouse sperm collection

The caudal epididymides were dissected from the *Cfap53* wildtype and knockout mice. Spermatozoa were squeezed out from the caudal epididymis and released in 1 ml phosphate buffered saline (PBS) for 30 min at 37°C under 5% CO_2_ for sperm counting and immunofluorescence experiments.

### Tissue collection and histological analysis

The testes and caudal epididymides from at least five *Cfap53* wildtype and five knockout mice were dissected immediately after euthanasia. All samples were immediately fixed in 4% (mass/vol) paraformaldehyde (PFA; Solarbio, P1110) for up to 24 hours, dehydrated in 70% (vol/vol) ethanol, and embedded in paraffin. For histological analysis, the 5 μm sections were mounted on glass slides and stained with H&E. For PAS staining, testes were fixed with Bouin’s fixatives (Polysciences). Slides were stained with PAS and H&E after deparaffinization, and the stages of the seminiferous epithelium cycle and spermatid development were determined.

### Immunofluorescence of the testicular germ cells

The mouse testis was immediately dissected and fixed with 2% paraformaldehyde in 0.05% PBST (PBS with 0.05% Triton X-100) at room temperature for 5 min. The fixed sample was placed on a slide glass and squashed by placing a cover slip on top and pressing down. The sample was immediately flash frozen in liquid nitrogen, and the slides were stored at −80°C for further immunofluorescence experiments (Wellard et al., 2018). After removing the coverslips, the slides were washed with PBS three times and then treated with 0.1% Triton X-100 for 10 min, rinsed three times in PBS, and blocked with 5% bovine serum albumin (Amresco, AP0027). The primary antibody was added to the sections and incubated at 4°C overnight, followed by incubation with the secondary antibody. The nuclei were stained with DAPI. The immunofluorescence images were taken immediately using an LSM 780 microscope (Zeiss) or SP8 microscope (Leica).

### Immunofluorescence in testes

The testes of *Cfap53* wildtype and knockout mice were fixed in 4% PFA at 4°C overnight, dehydrated in 70% (vol/vol) ethanol, and embedded in paraffin. For histological analysis, the 5 μm sections were mounted on glass slides, then deparaffinized and rehydrated, followed by antigen retrieval in 10 mM sodium citrate buffer (pH 6.0) for 15 min and washing three times in PBS, pH 7.4. After blocking with 5% BSA containing 0.1% Triton X-100, the primary antibodies were added to the sections and incubated at 4°C overnight, followed by incubation with the secondary antibody. The nuclei were stained with DAPI, and images were acquired on an SP8 microscope (Leica).

### Transmission electron microscopy

The testes from at least three *Cfap53* wildtype and knockout mice were dissected and pre-fixed in 2.5% (vol/vol) glutaraldehyde in 0.1 M cacodylate buffer at 4°C overnight. After washing in 0.1 M cacodylate buffer, samples were cut into small pieces of approximately 1 mm^3^, then immersed in 1% OsO_4_ for 1 hour at 4°C. Samples were dehydrated through a graded acetone series and embedded in resin for staining. Ultrathin sections were cut on an ultramicrotome and double stained with uranyl acetate and lead citrate, and images were acquired and analyzed using a JEM-1400 transmission electron microscope.

### Statistical analysis

All of the experiments were repeated at least three times, and the results are presented as the mean ± SD. The statistical significance of the differences between the mean values for the different genotypes was measured by the Student’s t-test with a paired, 2-tailed distribution. The data were considered significant for P < 0.05.

## Contributors

W.L. and H.B.L. designed the study and wrote the article. B.B.W. performed most of the experiments and analyzed the data. X.C.Y. performed the experiments and assisted in writing the manuscript. C.L., L.N.W., and X.H.L. performed some of the immunofluorescence experiments. All authors assisted in data collection, interpreted the data, provided critical input to the manuscript, and approved the final manuscript.

## Acknowledgements

This work was funded by the National Natural Science Foundation of China (grant 81925015), the Strategic Priority Research Program of the Chinese Academy of Sciences (grant XDA16020701) and Qilu Young Scholars Program of Shandong University. We would like to thank the State Key Laboratory of Membrane Biology, Institute of Zoology, Chinese Academy of Science, for our electron microscopy work, and we are grateful to Pengyan Xia for his help in preparing the electron microscopy sample.

---

**Supplementary Figure 1.**
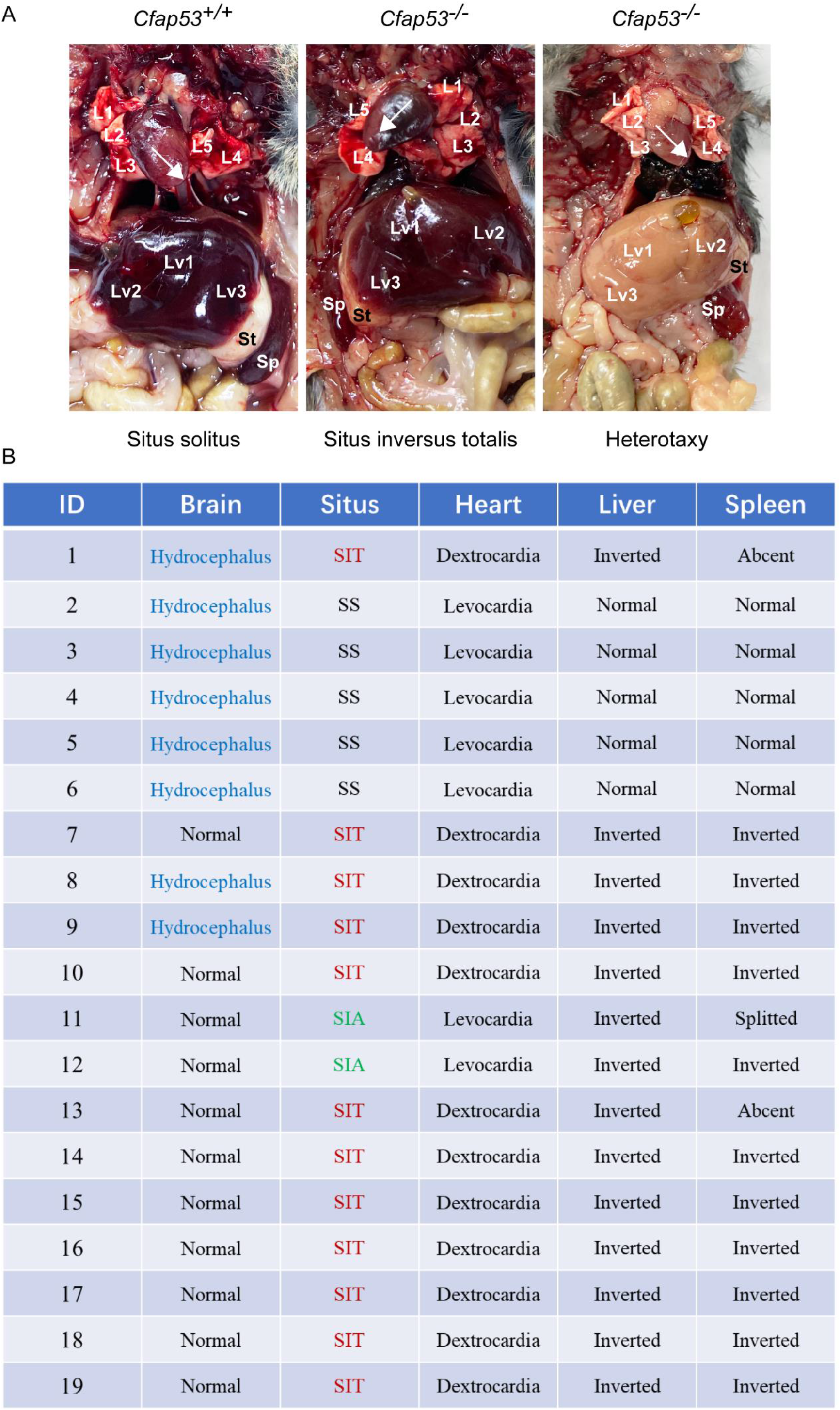
Left-right body asymmetry defects and hydrocephalus in *Cfap53*^−*/*–^ mice. (A) *Cfap53*^−*/*–^ mice presented with situs inversus totalis (SIT), situs inversus abdominalis (SIA), and situs solitus (SS). L1-5 (white numbers): lung lobes, Lv1-3 (white numbers): liver lobes, St: stomach, Sp: spleen. (B) Summary of the phenotypes detected in *Cfap53*^−*/*–^ mice.

**Supplementary Figure 2.**
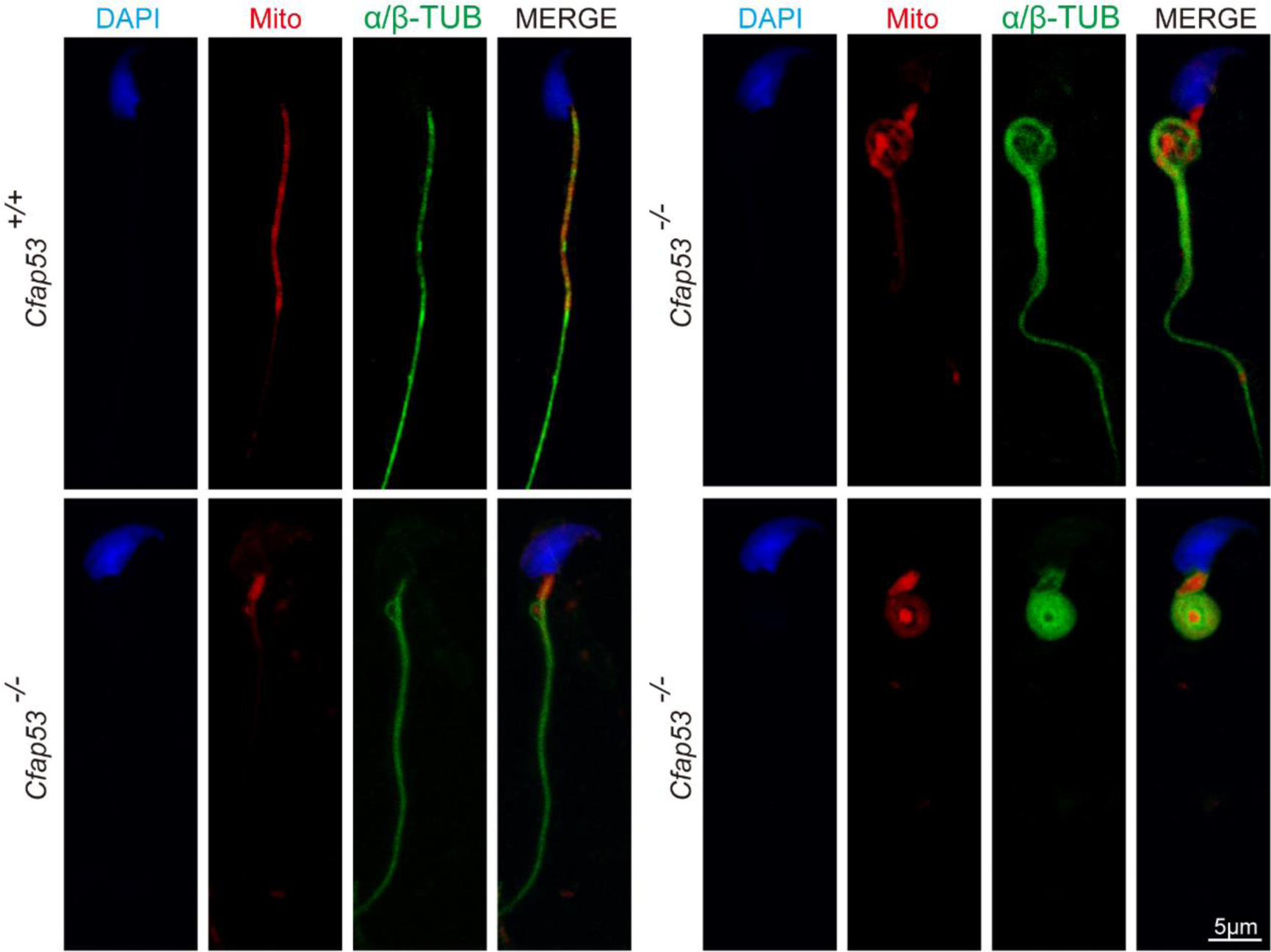
Mitochondrial sheath defects in *Cfap53*^−*/*–^ spermatozoa. The immunofluorescence analysis for α/β-tubulin (green) and MitoTracker (red) was performed in *Cfap53*^*+/+*^ and *Cfap53*^−*/*–^ spermatozoa. The nucleus was stained with DAPI (blue).

## References

Ben Khelifa, M., Coutton, C., Zouari, R., Karaouzène, T., Rendu, J., Bidart, M., Yassine, S., Pierre, V., Delaroche, J., Hennebicq, S., Grunwald, D., Escalier, D., Pernet-Gallay, K., Jouk, P. S., Thierry-Mieg, N., Touré, A., Arnoult, C., & Ray, P. F. (2014). Mutations in DNAH1, which encodes an inner arm heavy chain dynein, lead to male infertility from multiple morphological abnormalities of the sperm flagella. Am J Hum Genet 94(1), 95–104. https://doi.org/10.1016/j.ajhg.2013.11.017

Beurois, J., Martinez, G., Cazin, C., Kherraf, Z.-E., Amiri-Yekta, A., Thierry-Mieg, N., Bidart, M., Petre, G., Satre, V., & Brouillet, S. J. H. R. (2019). CFAP70 mutations lead to male infertility due to severe astheno-teratozoospermia. A case report 34(10), 2071–2079. https://doi.org/10.1093/humrep/dez166

Boivin, J., Bunting, L., Collins, J. A., & Nygren, K. G. (2007). International estimates of infertility prevalence and treatment-seeking: potential need and demand for infertility medical care. Hum Reprod 22(6), 1506–1512. https://doi.org/10.1093/humrep/dem046

Burgess, S. A., Walker, M. L., Sakakibara, H., Knight, P. J., & Oiwa, K. J. N. (2003). Dynein structure and power stroke. Nature 421(6924), 715–718. https://doi.org/10.1038/nature01377

Chemes, H. E., & Rawe, V. Y. (2010). The making of abnormal spermatozoa: cellular and molecular mechanisms underlying pathological spermiogenesis. Cell Tissue Res 341(3), 349–357. https://doi.org/10.1007/s00441-010-1007-3

Chen, H., Zhu, Y., Zhu, Z., Zhi, E., Lu, K., Wang, X., Liu, F., Li, Z., & Xia, W. (2018). Detection of heterozygous mutation in hook microtubule-tethering protein 1 in three patients with decapitated and decaudated spermatozoa syndrome. J Med Genet 55(3), 150–157. https://doi.org/10.1136/jmedgenet-2016-104404

Chien, A., Shih, S. M., Bower, R., Tritschler, D., Porter, M. E., & Yildiz, A. (2017). Dynamics of the IFT machinery at the ciliary tip. Elife, 6. https://doi.org/10.7554/eLife.28606

Coutton, C., Escoffier, J., Martinez, G., Arnoult, C., & Ray, P. F. (2015). Teratozoospermia: spotlight on the main genetic actors in the human. Hum Reprod Update 21(4), 455–485. https://doi.org/10.1093/humupd/dmv020

Dong FN, A.-Y. A., Martinez G, Saut A, Tek J, Stouvenel L, Lorès P, Karaouzène T, Thierry - Mieg N, Satre V, Brouillet S, Daneshipour A, Hosseini SH, Bonhivers M, Gourabi H, Dulioust E, Arnoult C, Touré A, Ray PF, Zhao H, Coutton C. (2018). Absence of CFAP69 c auses male Ninfertility due to multiple morphological abnormalities of the flagella in human and mouse. The American Journal of Human Genetics 102(4), 636–648. https://doi.org/10.1016/j.ajhg.2018.03.007

Escalier, D., & Touré, A. (2012). Morphological defects of sperm flagellum implicated in human male infertility. Med Sci (Paris) 28(5), 503–511. https://doi.org/10.1051/medsci/2012285015

He, X., Liu, C., Yang, X., Lv, M., Ni, X., Li, Q., Cheng, H., Liu, W., Tian, S., Wu, H., Gao, Y., Yang, C., Tan, Q., Cong, J., Tang, D., Zhang, J., Song, B., Zhong, Y., Li, H., Zhi, W., Mao, X., Fu, F., Ge, L., Shen, Q., Zhang, M., Saiyin, H., Jin, L., Xu, Y., Zhou, P., Wei, Z., Zhang, F., & Cao, Y. (2020). Bi-allelic Loss-of-function Variants in CFAP58 Cause Flagellar Axoneme and Mitochondrial Sheath Defects and Asthenoteratozoospermia in Humans and Mice. Am J Hum Genet 107(3), 514–526. https://doi.org/10.1016/j.ajhg.2020.07.010

Horowitz, E., Zhang, Z., Jones, B. H., Moss, S. B., Ho, C., Wood, J. R., Wang, X., Sammel, M. D., & Strauss, J. F., 3rd. (2005). Patterns of expression of sperm flagellar genes: early expression of genes encoding axonemal proteins during the spermatogenic cycle and shared features of promoters of genes encoding central apparatus proteins. Mol Hum Reprod 11(4), 307–317. https://doi.org/10.1093/molehr/gah163

Huang, T., Yin, Y., Liu, C., Li, M., Yu, X., Wang, X., Zhang, H., Muhammad, T., Gao, F., Li, W., Zijiang, C., Hongbin, L., & Jinlong, M. (2020). Absence of murine CFAP61 causes male infertility due to multiple morphological abnormalities of the flagella. Science Bulletin https://doi.org/10.1016

Ide, T., Twan, W. K., Lu, H., Ikawa, Y., Lim, L. X., Henninger, N., Nishimura, H., Takaoka, K., Narasimhan, V., Yan, X., Shiratori, H., Roy, S., & Hamada, H. (2020). CFAP53 regulates mammalian cilia-type motility patterns through differential localization and recruitment of axonemal dynein components. PLoS Genet 16(12), e1009232. https://doi.org/10.1371/journal.pgen.1009232

Kierszenbaum, A. L. (2001). Spermatid manchette: plugging proteins to zero into the sperm tail. Mol Reprod Dev 59(4), 347–349. https://doi.org/10.1002/mrd.1040

Kierszenbaum, A. L. (2002). Intramanchette transport (IMT): managing the making of the spermatid head, centrosome, and tail. Mol Reprod Dev 63(1), 1–4. https://doi.org/10.1002/mrd.10179

Kierszenbaum, A. L., Rivkin, E., & Tres, L. L. (2011). Cytoskeletal track selection during cargo transport in spermatids is relevant to male fertility. Spermatogenesis 1(3), 221–230. https://doi.org/10.4161/spmg.1.3.18018

Kierszenbaum, A. L., Rivkin, E., Tres, L. L., Yoder, B. K., Haycraft, C. J., Bornens, M., & Rios, R. M. (2011). GMAP210 and IFT88 are present in the spermatid golgi apparatus and participate in the development of the acrosome-acroplaxome complex, head-tail coupling apparatus and tail. Dev Dyn 240(3), 723–736. https://doi.org/10.1002/dvdy.22563

Kierszenbaum, A. L., & Tres, L. L. (2004). The acrosome-acroplaxome-manchette complex and the shaping of the spermatid head. Arch Histol Cytol 67(4), 271–284. https://doi.org/10.1679/aohc.67.271

Lehti, M. S., Kotaja, N., & Sironen, A. (2013). KIF3A is essential for sperm tail formation and manchette function. Mol Cell Endocrinol 377(1-2), 44–55. https://doi.org/10.1016/j.mce.2013.06.030

Lehti, M. S., & Sironen, A. (2016). Formation and function of the manchette and flagellum during spermatogenesis. Reproduction 151(4), R43–54. https://doi.org/10.1530/REP-15-0310

Lehti, M. S., & Sironen, A. (2017). Formation and function of sperm tail structures in association with sperm motility defects. Biol. Reprod 97(4), 522–536. https://doi.org/10.1093/biolre/iox096

Li, L., Sha, Y., Wang, X., Li, P., Wang, J., Kee, K., & Wang, B. (2017). Whole-exome sequencing identified a homozygous BRDT mutation in a patient with acephalic spermatozoa. Oncotarget 8(12), 19914–19922. https://doi.org/10.18632/oncotarget.15251

Li, W., He, X., Yang, S., Liu, C., Wu, H., Liu, W., Lv, M., Tang, D., Tan, J., & Tang, S. (2019). Biallelic mutations of CFAP251 cause sperm flagellar defects and human male infertility. Journal of human genetics 64(1), 49–54. https://doi.org/10.1038/s10038-018-0520-1

Li, W., Wu, H., Li, F., Tian, S., Kherraf, Z.-E., Zhang, J., Ni, X., Lv, M., Liu, C., & Tan, Q. (2020). Biallelic mutations in CFAP65 cause male infertility with multiple morphological abnormalities of the sperm flagella in humans and mice. J Med Genet 57(2), 89–95. https://doi.org/10.1136/jmedgenet-2019-106344

Li, Y., Sha, Y., Wang, X., Ding, L., Liu, W., Ji, Z., Mei, L., Huang, X., Lin, S., Kong, S., Lu, J., Qin, W., Zhang, X., Zhuang, J., Tang, Y., & Lu, Z. (2019). DNAH2 is a novel candidate gene associated with multiple morphological abnormalities of the sperm flagella. Clin Genet 95(5), 590–600. https://doi.org/10.1111/cge.13525

Liu, C., He, X., Liu, W., Yang, S., Wang, L., Li, W., Wu, H., Tang, S., Ni, X., Wang, J., Gao, Y., Tian, S., Zhang, L., Cong, J., Zhang, Z., Tan, Q., Zhang, J., Li, H., Zhong, Y., Lv, M., Li, J., Jin, L., Cao, Y., & Zhang, F. (2019). Bi-allelic Mutations in TTC29 Cause Male Subfertility with Asthenoteratospermia in Humans and Mice. Am J Hum Genet 105(6), 1168–1181. https://doi.org/10.1016/j.ajhg.2019.10.010

Liu, C., Lv, M., He, X., Zhu, Y., Amiri-Yekta, A., Li, W., Wu, H., Kherraf, Z.-E., Liu, W., & Zhang, J. (2020). Homozygous mutations in SPEF2 induce multiple morphological abnormalities of the sperm flagella and male infertility. Journal of Medical Genetics 57(1), 31–37. https://doi.org/10.1136/jmedgenet-2019-106011

Liu, C., Miyata, H., Gao, Y., Sha, Y., Tang, S., Xu, Z., Whitfield, M., Patrat, C., Wu, H., Dulioust, E., Tian, S., Shimada, K., Cong, J., Noda, T., Li, H., Morohoshi, A., Cazin, C., Kherraf, Z. E., Arnoult, C., Jin, L., He, X., Ray, P. F., Cao, Y., Touré, A., Zhang, F., & Ikawa, M. (2020). Bi-allelic DNAH8 Variants Lead to Multiple Morphological Abnormalities of the Sperm Flagella and Primary Male Infertility. Am J Hum Genet 107(2), 330–341. https://doi.org/10.1016/j.ajhg.2020.06.004

Liu, W., He, X., Yang, S., Zouari, R., Wang, J., Wu, H., Kherraf, Z. E., Liu, C., Coutton, C., Zhao, R., Tang, D., Tang, S., Lv, M., Fang, Y., Li, W., Li, H., Zhao, J., Wang, X., Zhao, S., Zhang, J., Arnoult, C., Jin, L., Zhang, Z., Ray, P. F., Cao, Y., & Zhang, F. (2019). Bi-allelic Mutations in TTC21A Induce Asthenoteratospermia in Humans and Mice. Am J Hum Genet 104(4), 738–748. https://doi.org/10.1016/j.ajhg.2019.02.020

Lv, M., Liu, W., Chi, W., Ni, X., Wang, J., Cheng, H., Li, W.-Y., Yang, S., Wu, H., & Zhang, J. (2020). Homozygous mutations in DZIP1 can induce asthenoteratospermia with severe MMAF. J Med Genet https://doi.org/10.1136/jmedgenet-2019-106479

Marszalek JR, R.-L. P., Roberts E, Chien KR, Goldstein LS. (1999). Situs inversus and embryonic ciliary morphogenesis defects in mouse mutants lacking the KIF3A subunit of kinesin-II. Proceedings of the National Academy of Sciences 96(9), 5043–5048. https://doi.org/10.1073/pnas.96.9.5043

Martinez, G., Kherraf, Z. E., Zouari, R., Fourati Ben Mustapha, S., Saut, A., Pernet-Gallay, K., Bertrand, A., Bidart, M., Hograindleur, J. P., Amiri-Yekta, A., Kharouf, M., Karaouzène, T., Thierry-Mieg, N., Dacheux-Deschamps, D., Satre, V., Bonhivers, M., Touré, A., Arnoult, C., Ray, P. F., & Coutton, C. (2018). Whole-exome sequencing identifies mutations in FSIP2 as a recurrent cause of multiple morphological abnormalities of the sperm flagella. Hum Reprod 33(10), 1973–1984. https://doi.org/10.1093/humrep/dey264

Mortimer, D. (2018). The functional anatomy of the human spermatozoon: relating ultrastructure and function. Mol. Hum. Reprod 24(12), 567–592. https://doi.org/10.1093/molehr/gay040

Narasimhan, V., Hjeij, R., Vij, S., Loges, N. T., Wallmeier, J., Koerner-Rettberg, C., Werner, C., Thamilselvam, S. K., Boey, A., Choksi, S. P., Pennekamp, P., Roy, S., & Omran, H. (2015). Mutations in CCDC11, which encodes a coiled-coil containing ciliary protein, causes situs inversus due to dysmotility of monocilia in the left-right organizer. Hum Mutat 36(3), 307–318. https://doi.org/10.1002/humu.22738

Noël, E. S., Momenah, T. S., Al -Dagriri, K., Al-Suwaid, A., Al-Shahrani, S., Jiang, H., Willekers, S., Oostveen, Y. Y., Chocron, S., Postma, A. V., Bhuiyan, Z. A., & Bakkers, J. (2016). A Zebrafish Loss-of-Function Model for Human CFAP53 Mutations Reveals Its Specific Role in Laterality Organ Function. Human Mutation 37(2), 194–200. https://doi.org/10.1002/humu.22928

Pasek, R. C., Malarkey, E., Berbari, N. F., Sharma, N., Kesterson, R. A., Tres, L. L., Kierszenbaum, A. L., & Yoder, B. K. (2016). Coiled-coil domain containing 42 (Ccdc42) is necessary for proper sperm development and male fertility in the mouse. Dev. Biol 412(2), 208–218. https://doi.org/10.1016/j.ydbio.2016.01.042

Perles, Z., Cinnamon, Y., Ta-Shma, A., Shaag, A., Einbinder, T., Rein, A. J. J. T., & Elpeleg, O. (2012). A human laterality disorder associated with recessive CCDC11 mutation. J Med Genet, 49(6) 386–390. https://doi.org/10.1136/jmedgenet-2011-100457

Ray, P. F., Toure, A., Metzler-Guillemain, C., Mitchell, M. J., Arnoult, C., & Coutton, C. (2017). Genetic abnormalities leading to qualitative defects of sperm morphology or function. Clin Genet 91(2), 217–232. https://doi.org/10.1111/cge.12905

San Agustin, J. T., Pazour, G. J., & Witman, G. B. (2015). Intraflagellar transport is essential for mammalian spermiogenesis but is absent in mature sperm. Mol Biol Cell 26(24), 4358–4372. https://doi.org/10.1091/mbc.E15-08-0578

Sha, Y., Liu, W., Wei, X., Zhu, X., Luo, X., Liang, L., & Guo, T. (2019). Biallelic mutations in Sperm flagellum 2 cause human multiple morphological abnormalities of the sperm flagella (MMAF) phenotype. Clin Genet 96(5), 385–393. https://doi.org/10.1111/cge.13602

Sha, Y., Wang, X., Yuan, J., Zhu, X., Su, Z., Zhang, X., Xu, X., & Wei, X. (2020). Loss-of-function mutations in centrosomal protein 112 is associated with human acephalic spermatozoa phenotype. Clin Genet 97(2), 321–328. https://doi.org/10.1111/cge.13662

Sha, Y., Wei, X., Ding, L., Mei, L., Huang, X., Lin, S., Su, Z., Kong, L., Zhang, Y., & Ji, Z. (2020). DNAH17 is associated with asthenozoospermia and multiple morphological abnormalities of sperm flagella. Ann Hum Genet 84(3), 271–279. https://doi.org/10.1111/ahg.12369

Sha, Y. W., Sha, Y. K., Ji, Z. Y., Mei, L. B., Ding, L., Zhang, Q., Qiu, P. P., Lin, S. B., Wang, X., Li, P., Xu, X., & Li, L. (2018). TSGA10 is a novel candidate gene associated with acephalic spermatozoa. Clin. Genet 93(4), 776–783. https://doi.org/10.1111/cge.13140

Shang, Y., Yan, J., Tang, W., Liu, C., Xiao, S., Guo, Y., Yuan, L., Chen, L., Jiang, H., Guo, X., Qiao, J., & Li, W. (2018). Mechanistic insights into acephalic spermatozoa syndrome-associated mutations in the human SUN5 gene. J Biol Chem 293(7), 2395–2407. https://doi.org/10.1074/jbc.RA117.000861

Silva, E., Betleja, E., John, E., Spear, P., Moresco, J. J., Zhang, S., Yates, J. R., Mitchell, B. J., & Mahjoub, M. R. (2016). Ccdc11 is a novel centriolar satellite protein essential for ciliogenesis and establishment of left-right asymmetry. Mol. Biol. Cell 27(1), 48–63. https://doi.org/10.1091/mbc.E15-07-0474

Sironen, A., Shoemark, A., Patel, M., Loebinger, M. R., & Mitchison, H. M. (2020). Sperm defects in primary ciliary dyskinesia and related causes of male infertility. Cell Mol Life Sci 77(11), 2029–2048. https://doi.org/10.1007/s00018-019-03389-7

Tang, S., Wang, X., Li, W., Yang, X., Li, Z., Liu, W., Li, C., Zhu, Z., Wang, L., & Wang, J. (2017). Biallelic mutations in CFAP43 and CFAP44 cause male infertility with multiple morphological abnormalities of the sperm flagella. The American Journal of Human Genetics 100(6), 854–864. https://doi.org/10.1016/j.ajhg.2017.04.012

Tapia Contreras, C., & Hoyer-Fender, S. (2019). CCDC42 Localizes to Manchette, HTCA and Tail and Interacts With ODF1 and ODF2 in the Formation of the Male Germ Cell Cytoskeleton. Front Cell Dev Biol 7, 151. https://doi.org/10.3389/fcell.2019.00151

Touré, A., Martinez, G., Kherraf, Z. E., Cazin, C., Beurois, J., Arnoult, C., Ray, P. F., & Coutton, C. (2020). The genetic architecture of morphological abnormalities of the sperm tail. Hum Genet https://doi.org/10.1007/s00439-020-02113-x

Tu, C., Nie, H., Meng, L., Yuan, S., He, W., Luo, A., Li, H., Li, W., Du, J., Lu, G., Lin, G., & Tan, YQ. (2019). Identification of DNAH6 mutations in infertile men with multiple morphological abnormalities of the sperm flagella. Sci Rep 9(1), 15864. https://doi.org/10.1038/s41598-019-52436-7

Tu, C., Wang, W., Hu, T., Lu, G., Lin, G., & Tan, YQ. (2020). Genetic underpinnings of asthenozoospermia. Best Pract Res Clin Endocrinol Metab 101472. https://doi.org/10.1016/j.beem.2020.101472

Turner, K. A., Rambhatla, A., Schon, S., Agarwal, A., Krawetz, S. A., Dupree, J. M., & Avidor-Reiss, T. (2020). Male Infertility is a Women’s Health Issue-Research and Clinical Evaluation of Male Infertility Is Needed. Cells, 9(4). https://doi.org/10.3390/cells9040990

Turner, R. M., Musse, M. P., Mandal, A., Klotz, K., Jayes, F. C., Herr, J. C., Gerton, G. L., Moss, S. B., & Chemes, H. E. (2001). Molecular genetic analysis of two human sperm fibrous sheath proteins, AKAP4 and AKAP3, in men with dysplasia of the fibrous sheath. J Androl 22(2), 302–315.

Tüttelmann, F., Ruckert, C., & Röpke, A. (2018). Disorders of spermatogenesis: Perspectives for novel genetic diagnostics after 20 years of unchanged routine. Med Genet 30(1), 12–20. https://doi.org/10.1007/s11825-018-0181-7

Wellard, S. R., Hopkins, J., & Jordan, P. W. (2018). A seminiferous tubule squash technique for the cytological analysis of spermatogenesis using the mouse model. JoVE(132) e56453. https://doi.org/10.3791/56453

Wu, B., Gao, H., Liu, C., & Li, W. (2020). The coupling apparatus of the sperm head and tail. Biol Reprod 102(5), 988–998. https://doi.org/10.1093/biolre/ioaa016

Zhang, Z., Li, W., Zhang, Y., Zhang, L., Teves, M. E., Liu, H., Strauss, J. F., 3rd, Pazour, G. J., Foster, J. A., Hess, R. A., & Zhang, Z. (2016). Intraflagellar transport protein IFT20 is essential for male fertility and spermiogenesis in mice. Mol Biol Cell 27(23), 3705–3716. https://doi.org/10.1091/mbc.E16-05-0318

Zhu, F., Liu, C., Wang, F., Yang, X., Zhang, J., Wu, H., Zhang, Z., He, X., Zhang, Z., Zhou, P., Wei, Z., Shang, Y., Wang, L., Zhang, R., Ouyang, Y. C., Sun, Q. Y., Cao, Y., & Li, W. (2018). Mutations in PMFBP1 Cause Acephalic Spermatozoa Syndrome. Am J Hum Genet 103(2), 188–199. https://doi.org/10.1016/j.ajhg.2018.06.010

Zhu, F., Wang, F., Yang, X., Zhang, J., Wu, H., Zhang, Z., Zhang, Z., He, X., Zhou, P., Wei, Z., Gecz, J., & Cao, Y. (2016). Biallelic SUN5 Mutations Cause Autosomal-Recessive Acephalic Spermatozoa Syndrome. Am J Hum Genet 99(4), 942–949. https://doi.org/10.1016/j.ajhg.2016.11.002

Zhu, X., Wang, J., Li, S., Lechtreck, K., & Pan, J. (2020). IFT54 directly interacts with kinesin-II and IFT dynein to regulate anterograde intraflagellar transport. Embo j, e105781. https://doi.org/10.15252/embj.2020105781

Zinaman, M. J., Brown, C. C., Selevan, S. G., & Clegg, E. D. (2000). Semen quality and human fertility: a prospective study with healthy couples. J Androl 21(1), 145–153.

